# Characterization of cells susceptible to SARS-COV-2 and methods for detection of neutralizing antibody by focus forming assay

**DOI:** 10.1101/2020.08.20.259838

**Authors:** E. Taylor Stone, Elizabeth Geerling, Tara L. Steffen, Mariah Hassert, Alexandria Dickson, Jacqueline F. Spencer, Karoly Toth, Richard J. DiPaolo, James D. Brien, Amelia K. Pinto

## Abstract

The SARS-CoV-2 outbreak and subsequent COVID-19 pandemic have highlighted the urgent need to determine what cells are susceptible to infection and for assays to detect and quantify SARS-CoV-2. Furthermore, the ongoing efforts for vaccine development have necessitated the development of rapid, high-throughput methods of quantifying infectious SARS-CoV-2, as well as the ability to screen human polyclonal sera samples for neutralizing antibodies against SARS-CoV-2. To this end, our lab has adapted focus forming assays for SARS-CoV-2 using Vero CCL-81 cells, referred to in this text as Vero WHO. Using the focus forming assay as the basis for screening cell susceptibility and to develop a focus reduction neutralization test. We have shown that this assay is a sensitive tool for determining SARS-CoV-2 neutralizing antibody titer in human, non-human primate, and mouse polyclonal sera following SARS-CoV-2 exposure. Additionally, we describe the viral growth kinetics of SARS-CoV-2 in a variety of different immortalized cell lines and demonstrate via human ACE2 and viral spike protein expression that these cell lines can support viral entry and replication.

## Introduction

Severe acute respiratory syndrome coronavirus 2 (SARS-CoV-2) is the etiological agent of the coronavirus disease 2019 (COVID-19) pandemic that rapidly traversed the globe beginning in December 2019, causing socio-economic upheaval and straining healthcare infrastructure worldwide. The long (~five day) incubation period before symptom onset, combined with aerosol-based transmission of SARS-CoV-2 has enabled the virus to ravage the immunologically naïve population and generate 7.2 million confirmed cases and over 400,000 deaths globally as of June 10th, 2020 [1, 2]. Since its isolation from a population of pneumonic patients in Wuhan within the Hubei province of China, the scientific community has mobilized a massive effort to develop sensitive and specific rapid-detection assays, as well as a vaccine that confers protection against SARS-CoV-2 infection, with several candidates currently under evaluation. However, there is an urgent need to identify susceptible cell lines for evaluation of potential antivirals and therapeutics for COVID-19 treatment. Understanding susceptibility of certain cell lines may also improve our understanding of SARS-CoV-2 viral pathogenesis. Furthermore, there is a need to develop rapid, high-throughput assays for the detection of infectious virus and neutralizing antibodies against SARS-CoV-2.

From phylogenetic analysis of early clinical isolates, it is known that SARS-CoV-2 is classified as a Group 2B betacoronavirus within the family *Coronavirinae* [3–6]. It is most closely related to severe acute respiratory syndrome coronavirus (SARS-CoV), the etiological agent of SARS, thought to have emerged from a zoonotic infection in China in the early 2000s [5, 7]. It is also more distantly related to Middle Eastern Respiratory Syndrome coronavirus (MERS-CoV) and other human common cold coronaviruses (HCoVs) [6]. Group 2B betacoronaviruses are enveloped viruses with a positive-sense RNA genome spanning 26-32kbp [8]. The protein coding genes are flanked by a 5’ cap and a 3’ polyadenylated tail. The genome is organized into structural and non-structural proteins. The 5’ end of the genome encodes the viral replicase in open reading frames (ORF) 1a and 1b, followed by a transcription cassette encoding subgenomic RNAs. SARS-CoV-2 encodes eight accessory proteins and four structural proteins: spike (S), envelope (E), membrane (M), and nucleocapsid (N), which are encoded by subgenomic RNAs located at the 3’ end of the genome and are necessary for the production of successful progeny virions [8, 9].

Much has been learned about the life cycle of SARS-CoV-2 in recent months. Both SARS-CoV-2 and SARS-CoV gain access to target cells by attachment of the viral spike protein to its cognate receptor, the protein angiotensin converting enzyme 2 (ACE2)[10–12]. In humans, ACE2 is highly expressed on epithelial cells throughout the oropharyngeal tract, lungs, and small intestine [13–15]. One subunit of the spike protein, S1, contains the receptor binding domain (RBD) and is responsible for binding to human ACE2 (hACE2) [10, 16]. A second spike protein subunit, S2, is highly conserved and coordinates with transmembrane serine protease 2 (TMPRSS2) to promote fusion with the host membrane [11, 17, 18]. While the spike-hACE2 interaction is a well-established determinant of susceptibility and host tropism for SARS-CoV-2, in order to combat the COVID-19 pandemic, scientists need to be armed with more information about the relationship between the expression of hACE2, cell permissivity, and viral titer, which has yet to be elucidated.

Scientists are also beginning to understand that the infectious dose and route of exposure may also play an important role in the development and severity of COVID-19. Particularly important when considering infectious dose and route of exposure are susceptible and permissible cell types. While it is known that the hACE2 receptor is required for entry, it is not well-understood which human cell types may be more permissive to SARS-CoV-2 infection, with particular uncertainty including if higher ACE2 expression coincides with increased risk for heightened infection. Furthermore, it is unclear which cell types may be suitable for the successful evaluation of antiviral and therapeutic drugs for *in vitro* screening before *in vivo* evaluation. Further characterization of permissible cell types is necessary to improve our understanding of SARS-CoV-2 pathogenesis and to develop therapeutic strategies for the treatment of COVID-19.

It has also been noted that following infection with SARS-CoV-2, people typically seroconvert 20 days following symptom onset, and there is increasing evidence suggesting that development of a neutralizing antibody response is a correlate of protection in patients recovered from COVID-19 [19–22]. The COVID-19 pandemic has highlighted the necessity for testing for neutralizing antibodies. As SARS-CoV-2 infection can have a range of manifestations from asymptomatic to fatal multiple organ failure [23–27], antibody testing and serological surveys are a critical tool for determining prior infection status and seroprevalence in a population. It is also the goal of many candidate SARS-CoV-2 vaccines to induce neutralizing antibodies targeting the viral spike protein, the major antigenic determinant of SARS-CoV-2 [28].

For understanding the antibody response, assays that measure neutralizing antibody titer are considered the gold standard. One such tool for evaluating neutralizing antibody response is a plaque/focus neutralization reduction test (PRNT/FRNT), which evaluates the ability of polyclonal sera samples to prevent or reduce infection of a cell monolayer in vitro. Previously, for SARS-CoV-2, only PRNT assays—which rely on the ability of virus to lyse infected cells and thus can take 48-96 hours to develop—have been used in the assessment of the neutralizing antibody response. It is not well known whether an FRNT—which uses an immunostaining protocol to detect virus and does not depend on cell lysis, and thus is often more rapid—is amenable for detection of SARS-CoV-2 neutralizing antibody.

In this study, we describe the growth kinetics of SARS-CoV-2 in multiple cell types and the methods our laboratory has used to optimize a SARS-CoV-2 focus forming assay (FFA) to improve sensitivity and specific detection. By characterizing the growth kinetics of SARS-CoV-2 on a variety of immortalized and primary cell lines, we have demonstrated which of these cell lines is susceptible to infection by SARS-CoV-2.We also demonstrate that the FFA can be adapted to measure the neutralization capacity of polyclonal sera in an FRNT. This high throughput FRNT assay can be applied to sera from both animal models of SARS-CoV-2 infection, as well as human SARS-CoV-2 infected patients, and can serve as a useful assay for describing the kinetics of the neutralizing antibody response to SARS-CoV-2. Additionally, we have compared the expression of ACE2 and SARS-CoV-2 spike on these cell lines to determine how spike expression correlates with susceptibility. The tools developed in this study have practical applications in both the basic science and translational approaches that will be critical in the ongoing effort to slow the COVID-19 pandemic.

## Results

### Cell types

Based on our previous work optimizing the FFA for WNV, [29] we proceeded to adapt the FFA for the detection of infectious SARS-CoV-2. Along with spot number, the spot size and border definition provide valuable information on possible differences in viral strains. As we have observed that the foci morphology, as well as spot number, can vary dramatically under different growth conditions, we sought to test different growth conditions and cell lines to determine the optimal conditions for SARS-CoV-2 viral titration. This goal was guided by previous studies that have suggested that the use of Vero cells from varying origins can impact viral titer [30, 31]. Using both Vero CCL-81 (ATCC^®^ CCL-81™, referred to in this text as Vero WHO) and Vero E6 (Vero 1008, ATCC^®^ CRL-1586™) cell lines, we determined if differences in foci number or size occurred to decide if one cell line was superior for titration by FFA (**Figure 1A–1C**). Although many laboratories utilize Vero E6 cells for viral titer measurements of SARS-CoV-2 [32, 33], in our laboratory, Vero E6 cells typically resulted in about two-fold lower foci formation relative to Vero WHO cells (**Figure 1A-C**). **Figure 1A** is a representative image of an FFA showing the viral titration on both the Vero E6 and WHOs for both ~50 FFU and ~200 FFU when identical numbers of cells are seeded per well. We noted that at identical higher dilutions of SARS-CoV-2 virus stocks, Vero WHO cells develop ~55 individual foci per well, whereas Vero E6 cells develop ~27 foci per well (**Figure 1A**). The same pattern was observed at lower dilutions of virus, with ~200 foci formation on Vero WHO cells yielding only ~100 foci on Vero E6 cells (**Figure 1A**) The quantification of this difference in the Vero WHO and Vero E6 cell type is shown in **Figure 1B.** Interestingly, when we compared this observation to the genome copy number by RT-qPCR, we noted that Vero E6 cells tended to produce significantly more virus across all 24, 48, and 72-hour timepoints (p = 0.0064) (**Figure 1C**). It is possible that the discrepancy between the FFA and qRT-PCR data could be due to Vero WHO cells producing fewer genome copies yet more infectious virus than E6 cells, while E6 cells sustain more viral replication but yield less infectious virus. While we did not see any differences in foci morphology between the two Vero cell lines we used, the significant difference in the number of foci observed between Vero E6 and Vero WHO (p < 0.0001) suggests that Vero WHO cells record higher viral titers than the Vero E6 cells for SARS-CoV-2 (**Figure 1B**).

**Figure 1.**
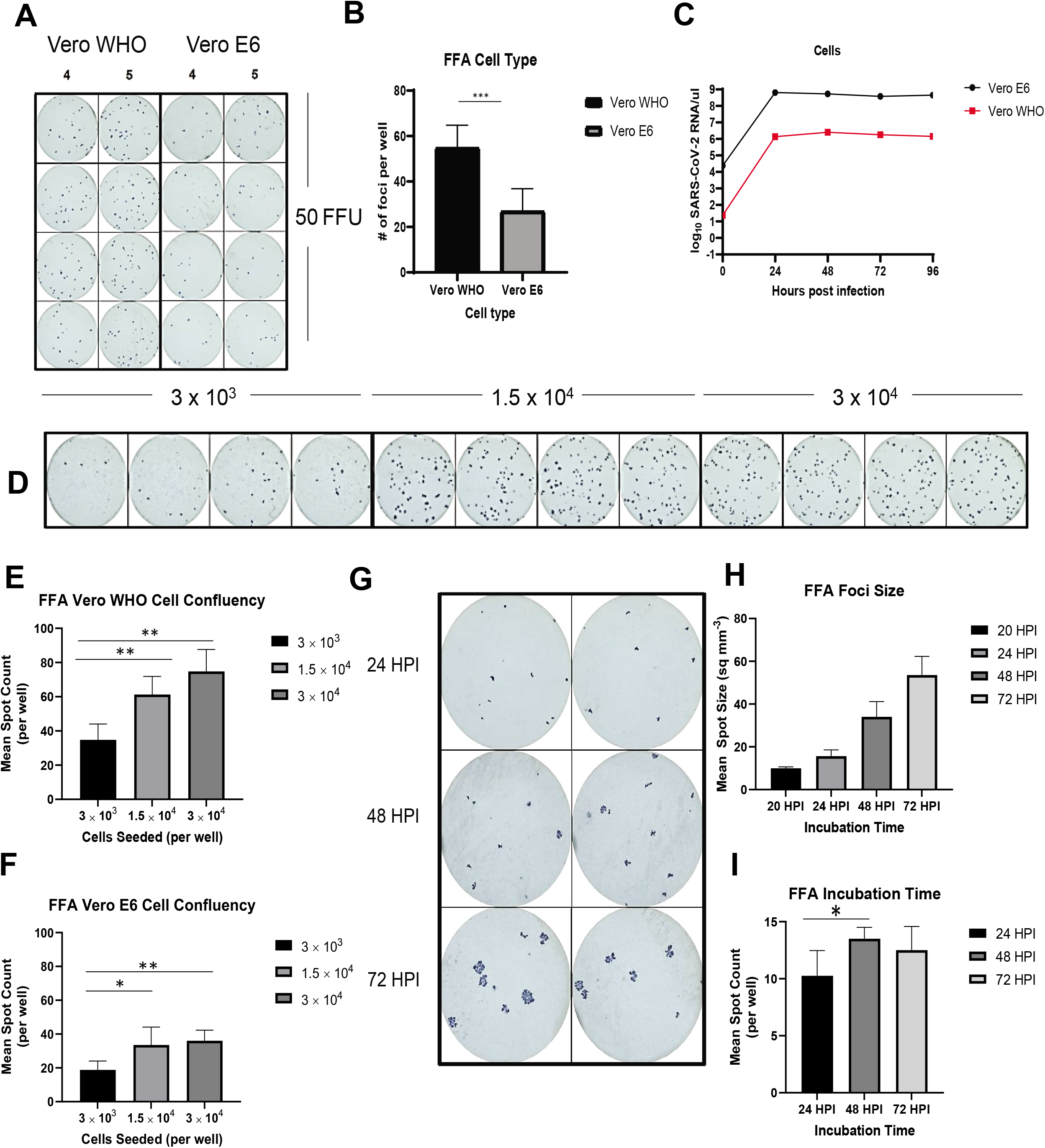
Detection of SARS-CoV-2 by focus forming assay (FFA). (A) Formation of foci using identical dilutions of virus on Vero WHO and Vero E6 cell types. Briefly, cells were seeded at 1.5 x 10^4^ cells/well in a 96-well plate one day prior to the assay. Identical dilutions of SARS-CoV-2 were allowed to infect cell monolayer for 1 hour at 37°C, 5% CO_2_ before overlay with 2% methylcellulose in 5% DMEM. Following a 24 hour incubation at 37°C, 5% CO_2_, cells were fixed in a solution of 5% paraformaldehyde diluted in 1 × PBS. Foci were visualized via immunostaining with polyclonal guinea pig anti-SARS-CoV-2 sera and goat anti-guinea pig conjugated HRP. KPL Trueblue Substrate was used for development of color and imaged the same day. (B) Quantification by RT-qPCR of SARS-CoV-2 genome copy number at 24 HPI for Vero WHO and Vero E6 cell types. (C) Representative image of difference in foci formation on Vero WHO and Vero E6 cell types following infection with identical dilutions of SARS-CoV-2. Differing numbers of Vero WHO (D) or Vero E6 (E) cells were seeded in 96-well plates and infected with identical dilutions of SARS-CoV-2 to determine the impact of on mean spot count per well. (F) Foci formation using identical dilutions of SARS-CoV-2 on different cell densities of Vero WHO cells. Impact of 20, 24, 48, and 72 hour incubation times on foci size (G) and formation (H) in a SARS-CoV-2 focus forming assay with 1.5 x 10^4^ cells seeded per well. (I) Representative images of identical dilutions of SARS-CoV-2 foci following 24, 48, and 72 hour incubation periods. Data displayed are the results of two independent experiments.

### Cell confluency

In order for SARS-CoV-2 to form distinct foci, it is critical to plate the optimal cell density. We examined the impact of cell density on foci formation for both Vero WHO and Vero E6 cells by plating identical dilutions of SARS-CoV-2 virus stocks on 96-well plates seeded with differing numbers of WHO or E6 cells (3 × 10^4^, 1.5 × 10^4^ or 3 × 10^4^ cells/well) one day prior to infection of the cell monolayer. At these concentrations, on the day of infection, 3×10^4^, 1.5×10^4^ and 3×10^4^ cells/well resulted in monolayers that were 70, 80 and 90 percent confluent respectively for both cell lines tested. **Figure 1D** is a representative image of the focus forming assay showing the foci formations arising from different cell concentrations plated for both the Vero E6 and WHOs. The quantification of the spot counts for Vero WHO cells is shown in **Figure 1E**. The quantification of the spot counts for Vero E6 cells is shown in **Figure 1F**. For both the Vero WHOs and E6 cells, we observed a significant increase in the number of foci formed when either 1.5 × 10^4^ (E6 p = 0.0480, WHO p = 0.0094) or 3 × 10^4^ (E6 p = 0.0057, WHO p = 0.0024) were seeded per well compared to 3 × 10^3^. However, there was no significant increase in foci formation when increasing the cell density from 1.5 × 10^4^ cells/well to 3 × 10^4^ cells/well. Thus, while the Vero E6 cells fostered lower foci numbers at each confluency as compared to the WHO cells, viral titers were not significantly different at 1.5 × 10^4^ cells/well or 3 × 10^4^ cells/well irrespective of whether the Vero WHO (**Figure 1E**) or Vero E6 (**Figure 1F**) cell line was seeded for the assay. We have previously tested higher cell densities for FFAs and have noted that cell concentrations higher than 3 × 10^4^ cells/well results in an overly confluent monolayer with more cells than can adhere to the wells, leading to highly variable titer information (data not shown). The results of these studies suggest that Vero WHO cells plated at either 1.5 × 10^4^ or 3 × 10^4^ cells/well was optimal for the viral FFA.

### Incubation times

Like plaque assays, the incubation time for the FFA is highly dependent on the viral replication cycle within the cells and the time required for infectious progeny to be released and spread to neighboring cells. While — unlike traditional plaque assays — the FFA is not dependent on viral lysis of infected cells, the development of visible spots is dependent on the time it takes for viral protein production to occur and for infectious virus to spread to neighboring susceptible cells. To determine the optimal time frame for infection of SARS-CoV-2 on a Vero WHO cell monolayer to form individual foci, we tested a variety of incubation times. In order to optimize these conditions, identical dilutions of SARS-CoV-2 virus stocks were added to infect Vero WHO cells seeded at a density of 1.5 × 10^4^ cells/well, and incubated for 20, 24, 48, or 72 hours post infection. **Figure 1G** shows representative images of identical dilutions of SARS-CoV-2 FFAs developed after 24, 48, or 72 hour incubation times, respectively. The mean spot size of foci at each timepoint is quantified in **Figure 1H.** The mean spot count per well at each time point is quantified in **Figure 1I.**

Altering the incubation time had the most dramatic impact on mean foci size amid all other parameters tested. While there were no significant differences in the size of foci formed between 20 hours post infection (HPI) and 24 HPI (p = 0.0632), we did observe the formation of significantly larger foci between the 24 and 48 HPI timepoints (p = 0.0031), as well as the 48 and 72 HPI timepoints (p = 0.0134) (**Figure 1H**). Interestingly, at the same virus dilution, we found that there were fewer spots between 24 HPI (mean spot number of 10) relative to the 48 HPI time point (p = 0.0369, mean spot number of 13.5) but not the 72 HPI time point (p = 0.1895, mean spot value 12.5). Similarly, we found that there were no significant differences in the number of foci formed between 48 and 72 HPI (p = 0.4198) (**Figure 1I**). However, we noted that this difference is within one standard deviation of the mean number of spots across all wells and was insignificant when determining titers. From this result we concluded that incubation times greater than 24 hours resulted in a slight but significant increase in spot number, while assays incubated for up to 72 did not alter the spot number but did increase the spot size. This increase in spot size but not number between 48 and 72 hours is a highly useful for the testing of anti-viral compounds which may require longer incubation times. In addition, larger spot size makes this assay more universally useful since laboratories without an automated machine can manually count spots, where the large size will improve readability of the assay.

The FFA relies on an immunostaining protocol of an infected cell monolayer in order to quantify infectious virus titer and is therefore dependent upon SARS-CoV-2-specific antibody binding. For this purpose, polyclonal guinea pig sera (BEI: NR-0361) raised against SARS-CoV-2 produces reproducible staining with minimal background. However, we have also used human monoclonal antibodies for this purpose with success. **Supplemental Figure 1A-1D** shows an optimized FFA immunostaining protocol using a human monoclonal antibody (Clone 2165, Leinco Technologies, Product No. LT1900) and goat anti-human IgG HRP (Invitrogen, Cat No. 62-8420) to detect infectious SARS-CoV-2 titer of virus stocks generated using Vero E6 and Huh7.5 cell lines.

### Viral replication in susceptible cell types

As efforts to understand replication and transmission of SARS-CoV-2 are underway and vaccine development moves forward, more information regarding permissive cell types for SARS-CoV-2 infection and replication is needed. To determine the permissivity of several different cell types to infection with SARS-CoV-2, we generated multistep growth curves for human, non-human primate, murine, hamster, and gastric adenocarcinoma cell lines. Each of these cell lines was infected at a MOI = 0.05 and cells or cell supernatants were collected aseptically, and total RNA was isolated. Cellular RNA was normalized to an internal RNaseP control for human and non-human primate cells and GAPDH for murine cells. Genome equivalents were determined using the Applied Biosystems TaqMan gene expression assay protocol for SARS-CoV-2 previously described [34].

To identify susceptible cell lines, we first assessed genome copy number in non-human primate African green monkey kidney epithelial cells (Vero WHO, Vero E6) as well as human hepatocytes (Huh7.5, Huh7) and lung epithelial cells (A549, CALU-3). **Figure 2A** shows the SARS-CoV-2 genome copy number for whole cells for human and non-human primate cell lines. **Figure 2B** shows the SARS-CoV-2 genome copy number for cell culture supernatants for human and non-human primate cell lines. In each cell line aside from Vero WHO and Huh7, genome equivalents within the cell peaked at 24 HPI and remained relatively constant for the duration of the experiment. SARS-CoV-2 genome equivalents in Vero WHO and Huh7 cells peaked at 48 HPI and remained relatively constant for the duration of the experiment. The Vero E6 cell line reached the highest titer at 6.44 × 10^8^ copies/μL at 24 HPI, with the Huh7.5 and CALU-3 cell lines reaching 1.1 × 10^6^ copies/μL and 5.0 × 10^5^ copies/μL, at the same time point, respectively. Vero WHO and Huh7 cell lines reached the highest titer, 1.3 × 10^6^ copies/μL and 1.1 × 10^6^ copies/μL, respectively, at 48 HPI. The A549 cells, although susceptible to infection, appeared to support little SARS-CoV-2 replication, reaching only 80 copies/μL at 24 HPI.

**Figure 2.**
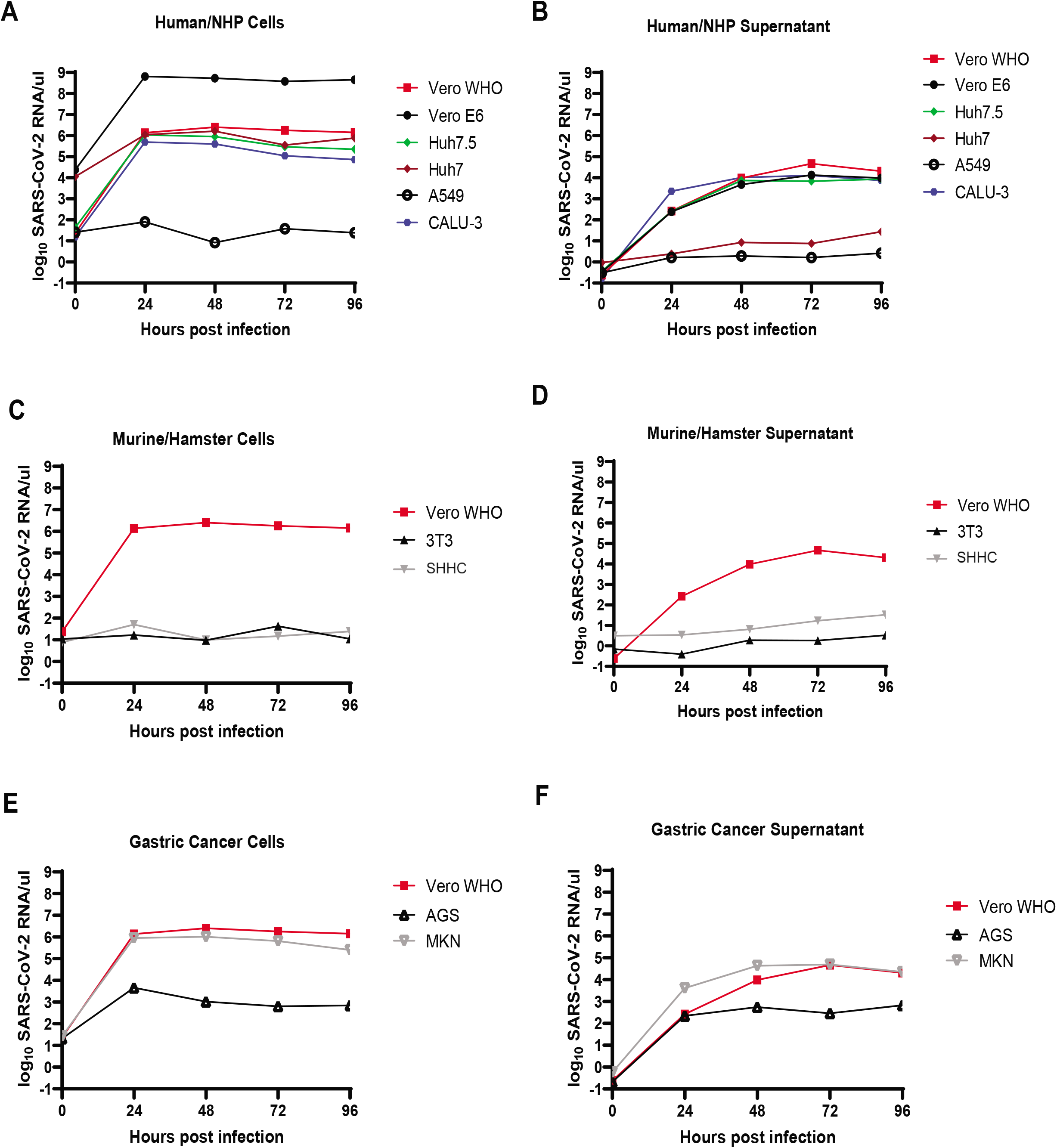
Viral replication and growth kinetics in immortalized cell lines. (A) SARS-CoV-2 genome copy number per μL as determined by RT-qPCR for human and non-human primate cell lines infected with SARS-CoV-2 at a MOI = 0.05 and sampled at 24-hour intervals. Total cellular RNA was extracted and normalized to a RNaseP control as well as a virus copy control. (B) SARS-CoV-2 genome copy number per μL as determined by RT-qPCR for the culture supernatants of human and non-human primate cell lines infected with SARS-CoV-2 at a MOI = 0.05 and sampled at 24-hour time intervals and normalized to a virus copy control. (C) SARS-CoV-2 genome copy number per μL as determined by RT-qPCR for rodent cell lines infected with SARS-CoV-2 at a MOI = 0.05 and sampled at 24-hour time intervals. Total cellular RNA was extracted using $kit$ according to manufacturer specifications and normalized to a GAPDH control as well as a virus copy control. Vero WHO cells are included for comparison. (D) SARS-CoV-2 genome copy number per μL as determined by RT-qPCR for the culture supernatants of rodent cell lines infected with SARS-CoV-2 at a MOI = 0.05 and sampled at 24-hour time intervals and normalized to a virus copy control. Vero WHO cell supernatant is included for comparison. (E) SARS-CoV-2 genome copy number per μL as determined by RT-qPCR for human gastric adenocarcinoma cell lines infected with SARS-CoV-2 at a MOI = 0.05 and sampled at 24-hour time intervals. Total cellular RNA was extracted using $kit$ according to manufacturer specifications and normalized to a GAPDH control as well as a virus copy control. Vero WHO cells are included for comparison. (F) SARS-CoV-2 genome copy number per μL as determined by RT-qPCR for the culture supernatants of human gastric adenocarcinoma cell lines infected with SARS-CoV-2 at a MOI = 0.05 and sampled at 24-hour time intervals and normalized to a virus copy control. Vero WHO cell supernatant is included for comparison.

**Figure 3.**
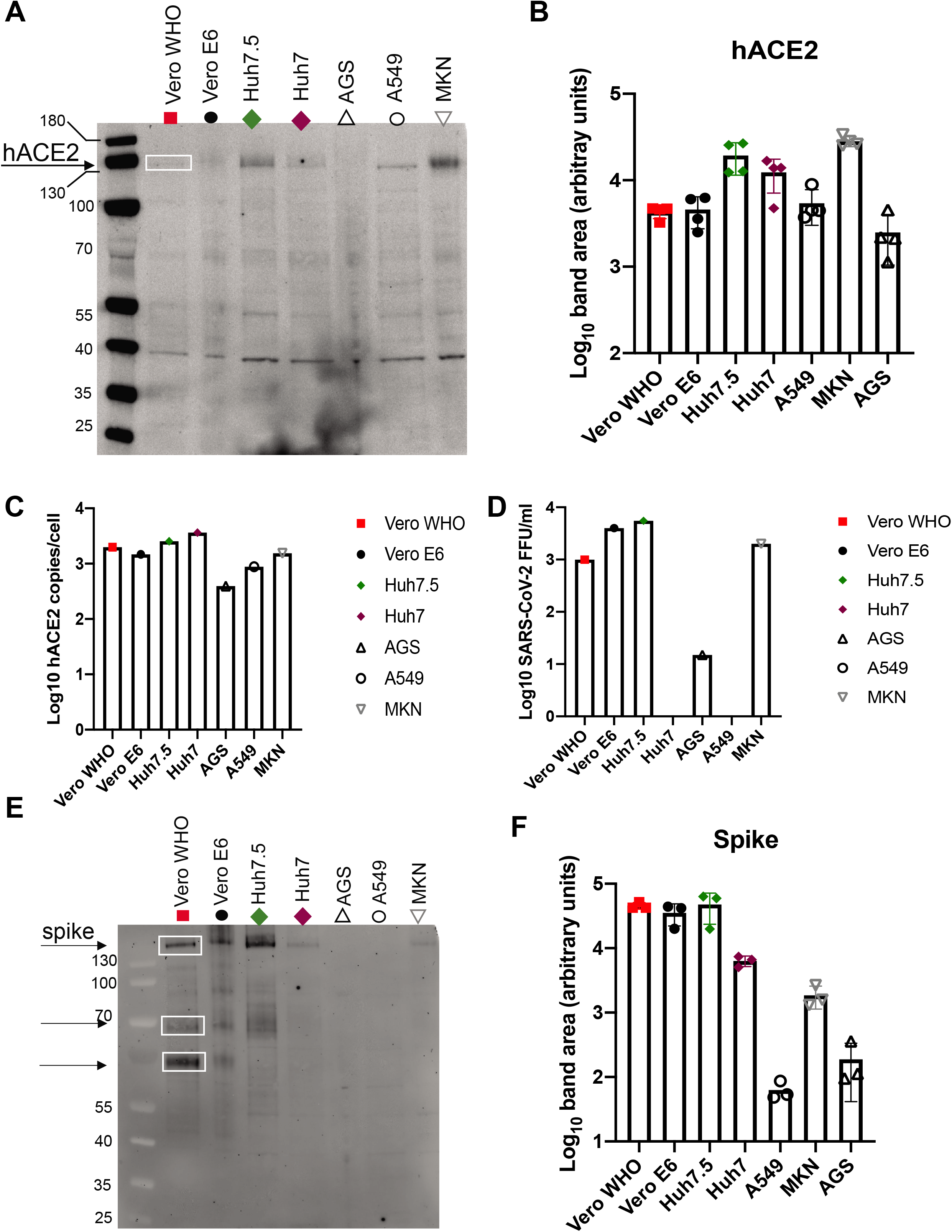
Cell lines expressing angiotension converting enzyme 2 (ACE2) and SARS-CoV-2 spike protein

We also measured genome copy number in the cell culture supernatant for all of the cell lines described (**Figure 2B**) For each cell line, viral RNA in the supernatant peaked at later timepoints, either 72 HPI (Vero WHO, 4.7× 10^4^ copies/μL; Vero E6, 1.4× 10^4^ copies/μL; CALU-3, 1.3× 10^4^ copies/μL) or 96 HPI (A549, 2.6 copies/μL; Huh7, 27 copies/μL; Huh7.5, 8.6 × 10^3^ copies/μL). Of these cell lines, the Vero WHO cells had the highest titers in the supernatant, while Vero E6, Huh7.5 and CALU-3 cells were comparable in terms of titer. As expected, A549 cells that did not contain high titers of cell-associated virus also did not contain high titers in the supernatant. Interestingly, while Huh7 cells support relatively high titers of cell-associated virus, they do not appear to yield high titers in the supernatant. These results suggest that Vero E6 cells are most permissible for SARS-CoV-2 replication among all tested cell types and would be the ideal choice for propagation. Vero WHO, Huh7.5, and CALU-3 cells are also permissible cell types for SARS-CoV-2 infection and replication, however A549 cells do not appear to be suitable for high levels of SARS-CoV-2 replication. Our results also suggest that Huh7 cells appear to be permissible for SARS-CoV-2 infection and replication, but do not appear to be suitable for egress into the cell culture supernatant.

Due to ongoing SARS-CoV-2 vaccine development efforts, there is an urgent need to develop and evaluate the susceptibility of small animal models to SARS-CoV-2 infection and COVID-19. Recent studies have suggested that rodents may be used for these purposes, as well as to study the adaptive immune response to SARS-CoV-2 infection [34–37]. To this end, we next sought to determine permissivity of rodent cell lines to SARS-CoV-2 infection, namely 3T3 and SHHC17 cell lines. **Figure 2C** shows the SARS-CoV-2 genome copy number for whole cells for murine and hamster cell lines, with Vero WHO cells included for comparison. **Figure 2D** shows the SARS-CoV-2 genome copy numbers for cell culture supernatants derived from murine and hamster cell lines, with Vero WHO supernatant included for comparison. From total cellular RNA, we detected only 42 copies/μL in 3T3 cells at 72 HPI, the time point at which the titer peaked. Similarly, with SHHC17 cells, we detected only 50 copies/μL at 24 HPI, the time point at which the titer peaked. In addition to total cellular RNA, we also examined supernatants from cell culture and predictably found peak titers of only 3 copies/μL in 3T3 cells and just 33 copies/μL in SHHC17 cells, both at 96 HPI. These results suggest that neither 3T3 nor SHHC17 cell lines are suitable for supporting SARS-CoV-2 replication or egress without further experimental manipulation.

Finally, because it is known that hACE2 is highly expressed by intestinal epithelial cells, we sought to examine the permissivity of human gastric adenocarcinoma cell lines to SARS-CoV-2 infection [15]. For this purpose, we used AGS and MKN cell lines, and examined viral genome copies associated with both total cellular RNA as well as the cell supernatant. **Figure 2E** shows the SARS-CoV-2 genome copy numbers for whole cells from gastric adenocarcinoma lines, with Vero WHO cells included for comparison. **Figure 2F** shows the SARS-CoV-2 genome copy numbers for cell culture supernatants from gastric adenocarcinoma lines, with Vero WHO supernatant included for comparison. We found that MKN cells yielded relatively high titers, with cell-associated virus peaking at 1.0× 10^6^ copies/μL at 24 HPI. Virus in the supernatant peaked at 5.0 × 10^4^ copies/μL at 72 HPI. AGS cells yielded lower titers, with cell-associated virus peaking at 4.6× 10^3^ copies/μL at 24 HPI and virus in the supernatant peaking at 6.7 × 10^2^ copies/μL 96 HPI. Interestingly, however, the titer in AGS cells appeared more variable compared to other time points, increasing at 48 and 96 HPI and dropping at 24 and 72 HPI. These results suggest that these gastric adenocarcinoma cell lines can support infection, replication and egress of SARS-CoV-2 as well as, or in some cases better than, Vero cell lines.

### Quantification of hACE2 and viral spike (S) protein by western blot

Given that our studies conducted to quantify SARS-CoV-2 viral genome copies in susceptible cell lines yielded results that highlighted highly permissive cell lines, like Vero WHO, while also distinguishing less permissive cell lines, like A549, we sought to analyze spike and hACE2 protein co-expression to determine if higher hACE2 expression correlated with higher susceptibility to SARS-Cov-2 infection.

### Quantification of neutralizing antibody by FRNT

One facet of our understanding of the current SARS-CoV-2 outbreak that is rapidly evolving is SARS-CoV-2 seroprevalence in the general population. At the same time, forming a better understanding of and ability to assess the kinetics of the neutralizing antibody response to SARS-CoV-2 could be essential in further vaccine and anti-viral development efforts. To this end, we adapted the SARS-CoV-2 FFA for the quantification of neutralizing antibody (nAb) titers in the form of an FRNT. This was accomplished by incubating serially diluted convalescent serum from SARS-CoV-2 infected individuals with a known quantity of infectious SARS-CoV-2 (~60 FFU) and measuring foci formation. Infection was normalized to a PBS control to reflect the percent neutralization of sera.

First, we sought to determine whether the FRNT could be used to detect a range of nAb concentrations in human samples. **Figure 4A** shows the neutralization curves for human sera samples, showing a decrease in virus neutralization as the serum is diluted out. These samples were collected from 4 human subjects at the University of Puerto Rico following a positive qPCR test for SARS-CoV-2. All subjects were in the convalescence period at the time of sample collection. These assays were performed using deidentified residual sera samples.

**Figure 4.**
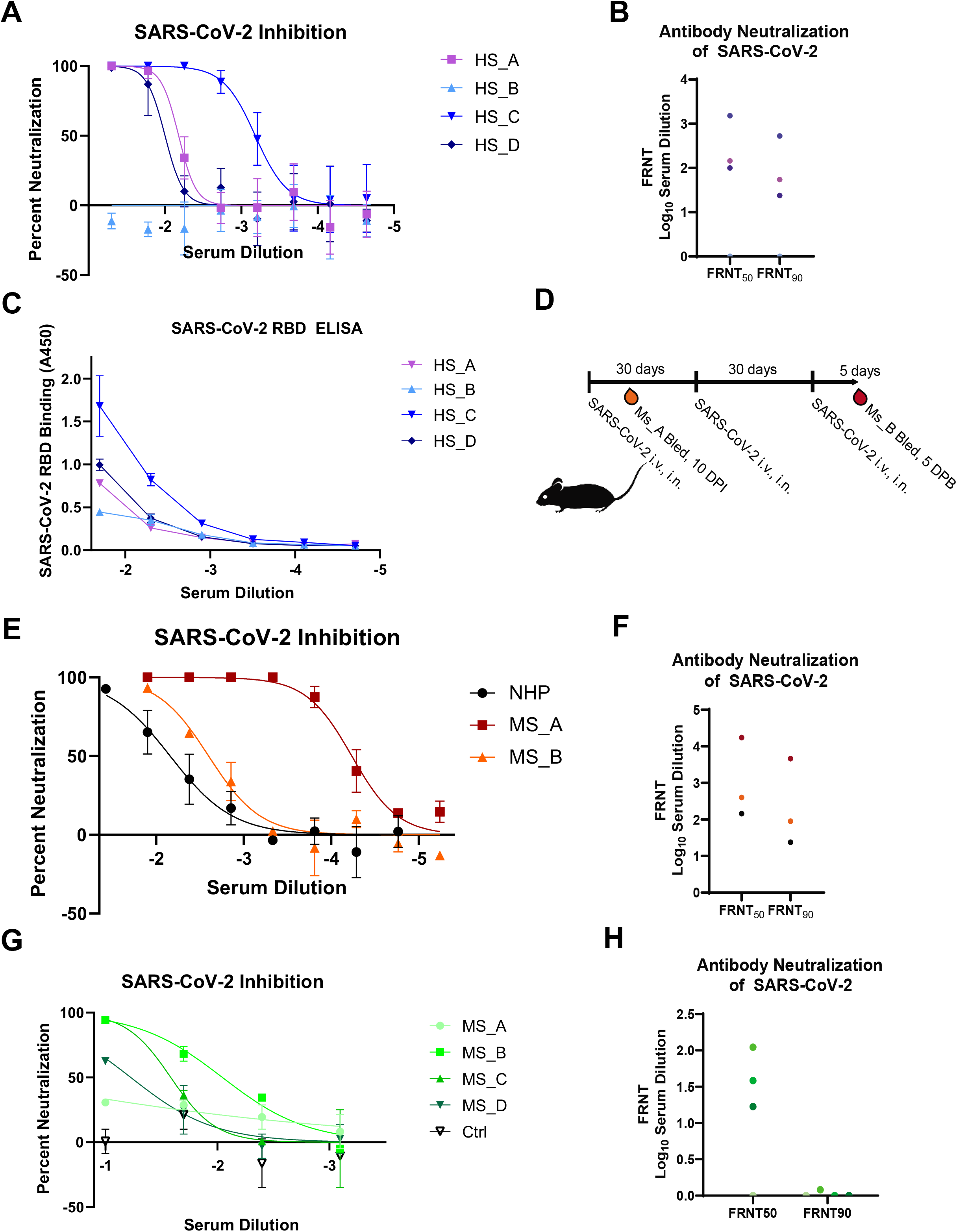
Quantification of neutralizing antibodies to SARS-CoV-2 in human and animal models of infection. (A) Inhibition of SARS-CoV-2 by three human subjects (HS_1, HS_2, and HS_3) >14 days post positive RT-qPCR test for SARS-CoV-2. The non-human primate sample (NHP_1) was taken from a pool of *Rhesus macaques* >14 days post inoculation with SARS-CoV-2 CDC isolate. The three mouse samples (MS_A, MS_B, MS_C, MS_D) were collected from IFNAR-/- mice on a C57BL/6 background 10 days post i.v. and i.n. inoculation with SARS-CoV-2. Neutralization capacity was measured by mixing serially diluted sera with a standardized amount of virus (~60 FFU) and allowed to incubate for 1 hour at 37°C, 5% CO_2_ before allowed to infect a cell monolayer as described in Figure 1. Inhibition of SARS-CoV-2 infection was normalized to a PBS control containing no sera. (B) The FRNT_50_ value as determined by focus reduction neutralization test. Dashed line indicates limit of detection. (C) Area under the curve (AUC) for binding to recombinant receptor binding domain (RBD) protein from SARS-CoV-2 for human, non-human primate, and mouse sera samples previously described.

Using the FRNT approach, we quantified the neutralizing antibody titer in the form of the reciprocal serum dilution required to neutralize 50% of virus, or the FRNT_50_ value. This assay can also be used to determine the FRNT_90_ value (i.e. required for 90% neutralization). The FRNT_50_ and FRNT_90_ values are reported in **Figure 4B**. The reciprocal serum dilutions required for 50% neutralization for the human samples (HS_A, HS_C and HS_D) are 2.161, 3.183, and 2.002, respectively. The reciprocal serum dilutions required for 90% neutralization for HS_A, HS_C and HS_D are 1.377, 2.725, and 1.739, respectively. For HS_B, the reciprocal serum dilution required to neutralize both 50% and 90% of the virus was below the lower limit of quantitation (LLOQ) for the assay.

In order to increase confidence that these nAbs were the result of recent SARS-CoV-2 infection rather than cross-reactivity with the four circulating human common cold coronaviruses, we performed an ELISA to examine binding of these sera samples to SARS-CoV-2 receptor-binding domain (RBD). **Figure 4C** shows the absorbance at 450 nm (A450) values indicating that sera from these subjects contain antibodies that can bind specifically to the receptor binding domain (RBD) of SARS-CoV-2.

Having confirmed that our assay can detect nAb to SARS-CoV-2 in human sera, we next sought to demonstrate that this assay is applicable for numerous sera sources including non-human primates and mice. To this end, we performed an FRNT with non-human primate (NHP) sera, which consisted of pooled sera samples from a group of *Rhesus macaques* in the convalescent phase following SARS-CoV-2 infection by multiple routes (BEI: NR-52401). **Figure 4E** shows the neutralization curve for this pool of NHP sera. Low but detectable nAb titers were present in this sample with an FRNT_50_ value of 2.161 and FRNT_90_ of 1.377, as depicted in **Figure 4F**.

Having shown that our assay can be utilized to quantify nAbs in NHP samples, we next sought to demonstrate that this assay could also be used for quantifying nAbs in small animal models such as mice. This also afforded us the opportunity to examine the nAb response both at an acute time point (MS_A) and following repeat challenge (MS_B). For these sera samples, two interferon receptor alpha (IFNAR1-/-) deficient mice were challenged with 5 × 10^4^ focus forming units (FFU) SARS-CoV-2. Sera was collected 10 days post infection (MS_A) or 65 days post initial challenge and 5 days following final challenge (MS_B) as described in **Figure 4D. Figure 4E** shows the neutralization curves for mice at both acute and amnestic time points. Low but detectable nAb titers were present for the acute time point (MS_A) with an FRNT_50_ value of 2.601 and FRNT_90_ of 1.953, as depicted in **Figure 4F**. Following repeated challenge with SARS-CoV-2, we were able to detect an increase in nAb titer, with FRNT_50_ of 4.238 and FRNT_90_ of 3.663.

Having demonstrated that our FRNT assay can be used to quantify nAb titers for human samples, non-human primates, and mice both in acute infection and memory responses, we next sought to determine whether this assay could be used to quantify nAb resulting from a subunit or DNA vaccine. To this end, we immunized C57BL/6 mice intramuscularly (i.m) with 50 μg of DNA encoding the SARS-CoV-2 spike (MS_C). A subset of these mice was boosted 21 days later (MS_B, MS_D) with 5 μg of DNA intramuscularly and sera collected 21 days following the boost. **Figure 4G** shows the neutralization curves for these mice immunized with DNA encoding the SARS-CoV-2 spike protein. No neutralization was detected for a control animal that received no immunization. From these data we were able to define the FRNT_50_ values for MS_B, MS_C, and MS_D, which are shown in **Figure 4H**, and are 2.045, 1.584, 1.227, respectively. However, the FRNT_90_ values were below the LLOQ for the assay, as well as the FRNT_50_ value for MS_A. These results suggest that the FRNT can be used to detect nAbs in sera resulting from immunizations, in addition to nAbs in human, NHP, and mouse sera resulting from SARS-CoV-2 infections.

## Discussion

To address the need for high-throughput, rapid quantification of infectious SARS-CoV-2, our group has developed a focus forming assay (FFA) for SARS-CoV-2 using Vero WHO cells. The strength of the FFA is the rapid visualization of individual foci forming from a single infectious unit or focus forming unit (FFU). The FFA for SARS-CoV-2 can be developed in as little as 24 hours, shorter relative to traditional plaque assays for human coronaviruses which can take 2-5 days [32, 38, 39]. The focus forming assay is also amenable to a 96-well plate format, allowing for assays to be scaled up or automated to handle large volumes of samples quickly relative to assays requiring plates with 24 wells or fewer. Automating the quantification of foci using equipment such as a CTL machine can also streamline the process of screening large numbers of samples. One potential disadvantage of the focus forming assay is the requirement of a SARS-CoV-2 specific antibody as the primary antibody for foci immunostaining. However, for our assays we have found that polyclonal guinea pig serum provides reproducible staining with minimal background when used at the appropriate concentrations, and numerous human monoclonal antibodies are now commercially available and suitable for this purpose [40–42].

In regard to the focus forming assay development, we initially hypothesized that the absence of in Vero E6 cells would make them more susceptible to SARS-CoV-2 infection and therefore a more sensitive choice of cell line for the focus forming assay. Surprisingly, we found that Vero WHO cells were more suited to foci formation. It is worth noting that other labs have shown that a higher titer and larger, clearer plaques result when Vero E6 cells are used in place of Vero WHO cells when performing plaque-assay based titrations with SARS-CoV-2 [32]. This may reflect differences between the Wuhan clinical isolate used in this study as opposed to the USA-WA1/2020 isolate or this may be an artifact of the focus forming assay. Because we find that by qPCR, genome copy number is typically highest in Vero E6 cells, we hypothesize that more defective or non-infectious virus results from replication in Vero E6 cells. Additionally, high levels of genome replication in Vero E6 may not correlate with ability to spread laterally in cell culture and form foci. The discrepancy in SARS-CoV-2 replication in these two cell lines warrants further study.

Our understanding of the impact of cell type on SARS-CoV-2 entry, replication, assembly, and egress is in its infancy. These gaps in our knowledge were recently made evident by the use of chloroquine and hydroxychloroquine—widely used anti-malarial drugs that create suboptimal conditions for pathogens by raising endosomal pH—in the treatment of COVID-19. These compounds were shown to be effective at preventing SARS-CoV-2 entry in Vero cells [43–45], but were shown to be ineffective as a post-exposure prophylactic in a randomized, double-blind, placebo-controlled clinical trial [46]. It has recently been shown that these compounds are not effective in inhibiting SARS-CoV-2 entry in human lung cell lines (CALU-3) [47]. It is hypothesized that this discrepancy across different cell lines and in patient populations exists because the expression of proteins required for SARS-CoV-2 entry, namely TMPSSR2 and Cathepsin L, is quite different between Vero cells and human lung cells [11, 48]. This scenario underscores the importance of more basic research into susceptible cell types and their suitability for *in vitro* drug screening, propagation, and quantification.

To advance our understanding of the SARS-CoV-2 life cycle in susceptible cell types, we generated multi-step growth curves for a variety of human, simian, and rodent cell types. In most cases, viral replication peaked at 24 HPI in susceptible cell lines and this cell-associated virus was maintained for the duration of the experiment. In many cases, however, the presence of virus in the supernatant did not peak until 72-96 HPI. In the context of the viral replication cycle, our data suggests that genome replication *in vitro* peaks after just 24 hours, however assembly and egress from infected cells may take as long as 72-96 HPI.

While there are conflicting reports concerning the suitability of Huh7 cells for SARS-CoV-2 studies [32, 33] we observed a striking discrepancy was between cell-associated virus within the total RNA and the virus detected within the cell supernatant. As much as 1.1 × 10^6^ copies/μL of cell-associated virus was detectable in RNA isolated from Huh7 cells, but virtually no detectable virus was found in the cell supernatant. This suggested to us that viral entry—and hence the production of cell-associated virus within the total RNA fraction—was independent of successful viral egress. This trend did not hold for RIG-I-deficient Huh 7.5 cells [49], suggesting that viral egress is interferon (IFN) sensitive. This observation is in accordance with previous studies describing SARS-CoV-2 sensitivity to type I IFNs [50, 51]. Further studies are warranted to determine what factors are necessary and sufficient for viral egress and could therefore serve as potential therapeutic targets.

Multiple groups have demonstrated success using golden Syrian hamsters to model SARS-CoV-2 infection, pathogenesis, and possibly transmission [35, 52–54], which may reflect differing susceptibility of hamster cell types based on anatomical location of the isolated cells.

We did not observe a strong correlation between ACE2 and viral spike protein levels, nor did we see a strong relationship between viral genome copy and ACE2 mRNA level. Our results suggest that host cell susceptibility to SARS-CoV-2 infection is more complicated than ACE2 expression alone, thus warranting further investigation.

### Quantification of neutralizing antibody by FRNT

We have showed by ELISA that convalescent sera sourced from human, non-human primate, and mice infected with SARS-CoV-2 can bind to the RBD of SARS-CoV-2. While the S2 subunit of the SARS-CoV-2 spike protein is highly conserved among betacoronaviruses, previous studies have showed that the RBD within the spike protein of SARS-CoV-2 is unique [16, 55]. In serological studies of SARS-CoV-2, the presence of antibodies binding SARS-CoV-2 RBD is considered the most sensitive and specific indicator of previous SARS-CoV-2 exposure. These results increase our confidence that within these polyclonal sera samples are neutralizing antibodies that are specific to SARS-CoV-2, rather than cross-reactivity due prior coronavirus exposure, as has been called into question by some [28, 55, 56].

As other labs have noted [57] we observed that binding to SARS-CoV-2 RBD appears to correlate with neutralization capacity, as human samples with a high AUC for RBD binding by ELISA also had lower EC50 values, indicating that low concentrations of sera from these patients were sufficient to neutralize 50% of a standardized amount of virus. We have showed that each of these samples can effectively neutralize SARS-CoV-2 *in vitro* and that neutralization can be measured via FRNT.

### Conclusions

In these studies, we have highlighted the utility of the FFA and FRNT assays in the characterization of the antibody response to SARS-CoV-2. These assays are amenable to adaptation in a clinical setting to study human disease and in animal model-based approaches of studying SARS-CoV-2 infection. We have also described the replication and viral growth kinetics of SARS-CoV-2 in a variety of immortalized cell lines and showed that these cell lines can support genome replication and viral protein production. Our findings underscore the need for further research into the mechanisms of viral life cycle, immune evasion and basic science while also emphasizing the need for translational research to aid in vaccine development and antiviral drug discovery. The tools described here have practical applications in both the basic science and translational approaches that will be critical in the ongoing effort to slow the COVID-19 pandemic.

## Methods

### Ethics statement

All animal studies were conducted in accordance with the Guide for Care and Use of Laboratory Animals of the National Institutes of Health and approved by the Saint Louis University Animal Care and Use Committee (IACUC; protocol 2771).

### Viruses and Cells

SARS-CoV-2 stocks were derived from SARS-CoV-2 (Isolate USA-WA1/2020) from BEI (Catalog #: NR-52281). Virus was passaged two times in Vero E6 cells (ATCC^®^ CRL-1586™) before clarification by centrifugation (3000 rpm for 30 min) and storage at −80°C until further use. For determination of viral titer via focus forming assay, Vero WHO cells (ATCC^®^ CCL-81™) were used. Unless otherwise specified, all cells were cultured in Dulbecco’s Modified Eagle Medium (Sigma-D5796-500ML) containing 1% HEPES (Sigma-H3537-100ML) and 5% FBS (Sigma-F0926) at 37°C, 5% CO_2_ (5% DMEM). AGS and MKN cells were gifted by R. J. DiPaolo. SHHC17 cells were gifted by K. Toth.

### Focus forming assay

One day prior to the assay, Vero WHO cells were plated in a 96 well flat bottom tissue culture treated plate. A cell density of ~1.5 x 10^4^ cells per well is ideal, but target confluency is 90-95% on the day of the assay. For determination of titer of a sample or virus stock, serial 10-fold dilutions of sample were made in a 96-well round bottom plate containing 5% DMEM. Media was then removed from the 96-well flat bottom plate containing the Vero WHO cell monolayer and replaced with 100μL per well of diluted samples. This plate containing sample dilution on the cell monolayer was placed in an incubator with 37°C, 5% CO_2_ for 1 hour. A solution of 2% methylcellulose (Sigma-M0512-250G) in 1 × PBS was made in advance of the assay and stored at 4°C until ready to use. On the day of the assay and during the one-hour infection period, 2% methylcellulose was diluted 1:1 in 5% DMEM and placed on a rocker to mix. The 1% methylcellulose-media mixture (hereby referred to as overlay media) was stored at room temperature until ready to use. After the one-hour infection period, the 96 well plate containing sample dilution and cell monolayer was removed from incubator. Overlay was added to the plate by adding 125 μL of overlay media to each well. This step reduces the uncontrolled spread of virus throughout the monolayer on the well, making it difficult to distinguish individual foci. After the addition of overlay media, the plate was returned to an incubator with 37°C, 5% CO_2_ for 24 hours. Following the 24 hour incubation, the 96-well plate was fixed in a solution of 5% paraformaldehyde (PFA) diluted in tissue culture grade 1 × PBS. An electron-microscopy grade paraformaldehyde (Fisher, Cat No: EMS-15713-S) is recommended to preserve the integrity of the viral proteins and improve the efficiency of the downstream immunostaining and detection steps. The plate was removed from the incubator and the media containing the overlay and sample was aspirated off. One wash with 150μL of 1 × PBS per well was performed, taking care not to disrupt the cell monolayer. 50μL per well of 5% PFA in PBS was added for the fixing step. With the 5% PFA still in the plate, the plate was submerged in a bath of 5% Formalin buffered phosphate (Fisher: SF100-4) in 1 × PBS for 15 minutes at room temperature. After 15 minutes, the plate was removed from the formaldehyde bath and the 5% PFA removed from the monolayer. One wash with 100μL of 1 × PBS (tissue culture grade) per well was performed. The plate was submerged in a bath of 1 × PBS to rinse and removed from BSL-3 containment. Foci were visualized by an immunostaining protocol. The 96-well plate was first washed twice with 150μL per well of FFA Wash buffer (1 × PBS, 0.05% Triton X-100). The primary antibody consisted of polyclonal anti-SARS-CoV-2 guinea pig sera (BEI: NR-0361) and was diluted 1:15,000 with FFA Staining Buffer (1 × PBS, 1mg/ml saponin (Sigma: 47036)).Then, 50μL per well of primary antibody was allowed to incubate for 2 hours at room temperature or 4°C overnight. The 96-well plate was then washed three times with 150μL per well of FFA Wash Buffer. The secondary antibody consisted of goat anti-mouse conjugated horseradish peroxidase (Sigma: A-7289) diluted 1:5,000 in FFA Staining Buffer. Similarly, 50μL per well of secondary antibody was allowed to incubate for 2 hours at room temperature or 4°C overnight. The plate was washed three times with 150uL per well of FFA wash buffer. Finally, 50μL per well KPL Trueblue HRP substrate was added to each well and allowed to develop in the dark for 10-15 minutes, or until blue foci are visible. The reaction was then quenched by two washes with Millipore water and imaged immediately thereafter with a CTL machine to quantify foci.

### Focus reduction neutralization test

Briefly, sera samples were diluted 1:40 in 5% DMEM and added to the topmost row of a round bottom 96-well plate. Sera samples were then serially diluted 1:3 down the remainder of the plate in 5% DMEM. An equal volume of SARS-CoV-2 diluted to ~600 FFU/mL (~60 FFU/100μL) was then added to the serially diluted sample, mixed thoroughly, and allowed to incubate at 37°C for 1 hour. Then 100μL of SARS-CoV-2+sera mixture was transferred to a Vero WHO cell monolayer (as described in the focus forming assay). From this point, the assay was as described in the focus forming assay section.

### Enzyme-linked immunosorbent assay (ELISA)

Binding of human polyclonal sera to recombinant SARS-CoV2 proteins was determined by ELISA. A 1ug/mL mixture of 50μL per well containing recombinant protein in carbonate buffer (0.1M Na2CO3 0.1M NaHCO3 pH 9.3) was used to coat MaxiSorp (ThermoFisher) 96-well plates overnight at 4°C. Plates were blocked with blocking buffer (PBS + 5%BSA + 0.5% Tween) for 2 hours at room temperature the following day and washed four times with wash buffer. Polyclonal sera was serially diluted in blocking buffer prior to plating. Sera was allowed to incubate for 1 hour at room temperature and washed four times with wash buffer. Following the one hour incubation, goat-anti-human IgG HRP (Sigma) conjugated secondary (1:5000) was added and allowed to incubate for 1 hour at room temperature. The plate was washed again four times with wash buffer and the ELISA was visualized with 100μL per well of TMB enhanced substrate (Neogen Diagnostics) and allowed to develop in the dark for 15 minutes. A solution of 1N HCl was used to quench the reaction and the plate was read for an optical density of 450 nanometers on a BioTek Epoch plate reader. The total peak area under the curve (AUC) was calculated using GraphPad Prism 8.

### Isolation of RNA from cell culture and culture supernatants

RNA was isolated from cell culture and supernatant using an Invitrogen Purelink RNA mini kit according to the manufacturer’s instructions.

### RT-qPCR

hACE2 expression was measured by qRT-PCR using Taqman primer and probe sets from IDT (assay ID Hs.PT.58.27645939). SARS-CoV-2 viral burden was measured by qRT-PCR using Taqman primer and probe sets from IDT with the following sequences: Forward 5’ GAC CCC AAA ATC AGC GAA AT 3’, Reverse 5’ TCT GGT TAC TGC CAG TTG AAT CTG 3’, Probe 5’ ACC CCG CAT TAC GTT TGG TGG ACC 3’. Synthesized hACE2 RNA was used as a copy control to quantify the number of hACE2 molecules present in each sample. Similarly, a SARS-CoV-2 copy number control (available from BEI) was used to quantify SARS-CoV-2 genomes.

## Acknowledgements

Financial support-This work was supported by Saint Louis University COVID-19 research Seed Funding to awarded to AKP and awarded to JDB.

## Author Contributions

ETS, EG, JDB, and AKP conceptualized the work and wrote and edited the manuscript. Viral stocks were grown and quantified by ETS. Focus forming assay was optimized by ETS. Growth curves, cell lysates and RNA were generated by EG. RT-qPCR and Western blotting were performed by EG. SARS-CoV-2 RBD ELISAs were completed by TLS. SARS-CoV-2 infections and harvests were performed by MH and JDB. Focus Reduction Neutralization Tests were completed by ETS. All authors reviewed and approved the final version of the manuscript.

