## Supplementary figures and images for "Characterization of cells susceptible to SARS-COV-2 and methods for detection of neutralizing antibody by focus forming assay"

### Supplemental Figure 1

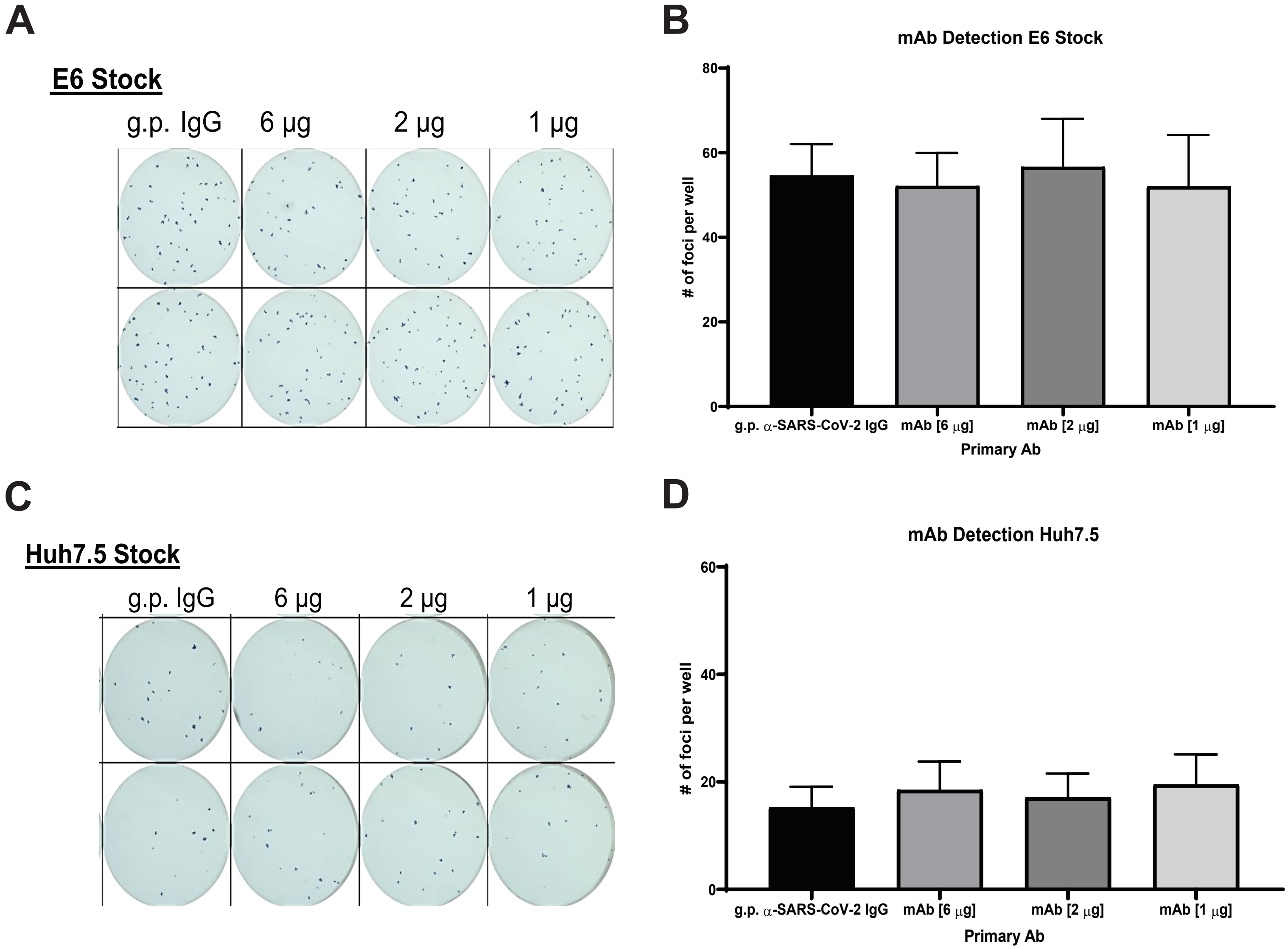
